## Supplementary Information for "The CASwitch: a synthetic biology solution for high-performance inducible gene expression systems in biotechnology"

a

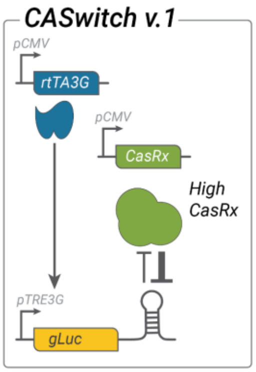


b

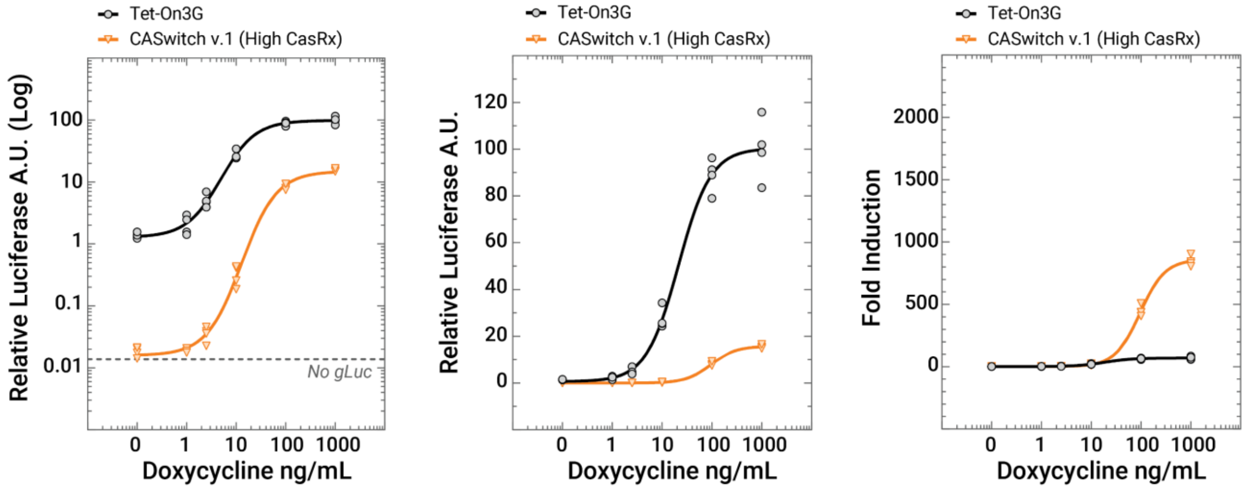


**Supplementary Figure 1. Characterization of the CASwitch v.1 system at higher relative concentrations of the CasRx endoribonuclease. (a)** CASwitch v.1 inducible gene expression system. The rtTA3G and the CasRx are constitutively expressed from constitutive pCMV promoter, while the gLuc with the Direct Repeat (DR) is placed downstream of the pTRE3G promoter. Cells were transfected with 5 times higher amount of the plasmid econding CasRx as compared to those used in Figure 2 d-f. **(b)** Experimental validation of CASwitch v.1 at relative higher concentration of CasRx (High CasRx, orange triangles) and comparison with the state-of-the-art Tet-On3G gene expression system (black circles) at the indicated concentrations of doxycycline. n=4 biological replicates are shown. Relative Luciferase A.U. of each data point is shown as the percentage of the mean of Luciferase A.U. value of the Tet-On3G system at 1000 ng/mL of doxycycline and plotted in a log-scale, linear-scale, and in as fold-induction values computed as the ratio between Relative Luciferase A.U of each data point and the mean of Relative Luciferase A.U in the absence of doxycycline. “No gLuc” represents the average Luciferase A.U. values of n=4 biological replicates of cells that were not transfected with the pTRE3G-gLuc-DR construct but only the pCMV-RedFireFly plasmid as transfection control, and shows the instrument background signal.

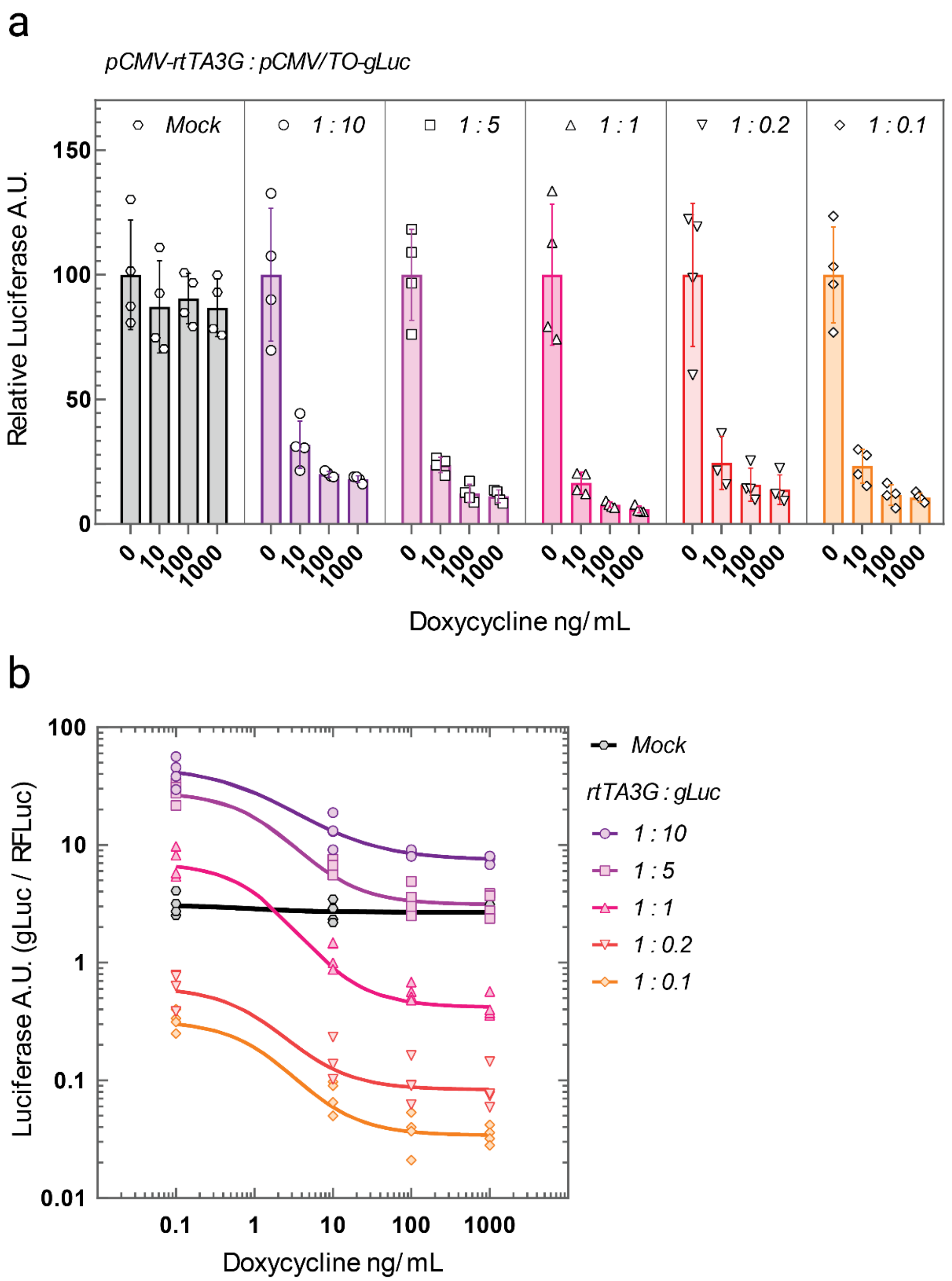


**Supplementary Figure 2. pCMV/TO promoter characterization.** Experimental validation of pCMV/TO transcriptional inhibitory function. HEK293T cells are transfected with two plasmids, one for the rtTA3G inducible transcriptional factor construct and the other the gLuc construct driven by the pCMV/TO promoter at the indicated relative concentrations. Luciferase expression was evaluated by dual-luminescence measurements at increasing amount of doxycycline. (a) Luciferase expression is plotted as Relative Luciferase A.U. which is the ratio between the Luciferase A.U. value of each data point and the mean of Luciferase A.U. values of the pCMV/TO-gLuc in the absence of doxycycline. Error bars correspond to the standard deviation for n = 4 biological replicates (b) Luciferase A.U. values calculated as the ratio between the luminescence value of gLuc and Red Firefly Luciferase of each data point.

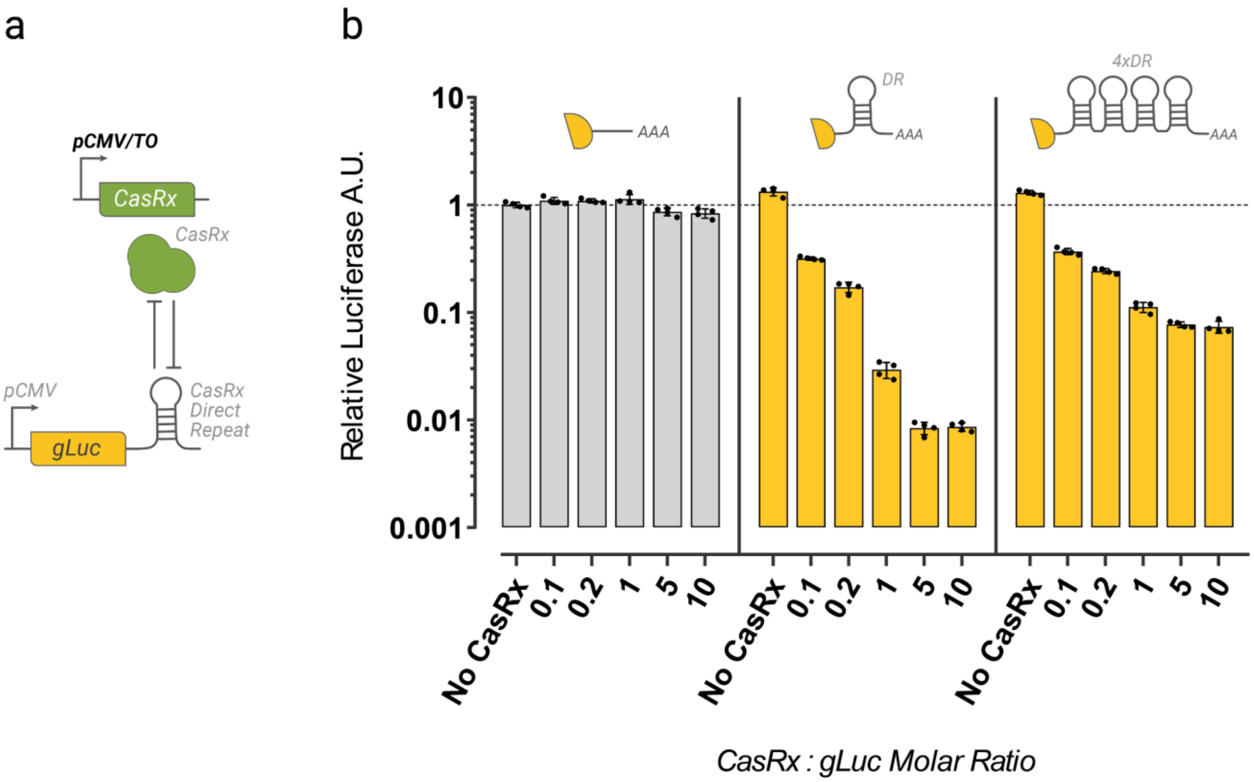


**Supplementary Figure 3. Mutual inhibition module featuring pCMV/TO-CasRx (a)** Schematic of the experimental implementation of the mutual inhibition module featuring a pCMV/TO-driven CasRx. Cells are transfected with two plasmids at the indicated relative concentrations, one for the pCMV/TO CasRx construct and the other for the gLuc construct harboring no, one and four repetitions of the Direct Repeat (DR) in the 3'Untranslated Region (3'UTR) of the gLuc reporter gene. **(b)** Experimental verification of the modified MI module. gLuc expression was evaluated by dual luminescence measurements resulting in the Luciferase A.U. value, calculated as the ratio between gLuc and Red Firefly Luciferase luminescence for each data point. Results are plotted as Relative Luciferase A.U. representing the ratio between the Luciferase A.U. value of each data point and the mean of Luciferase A.U. values of the gLuc without DR in the absence of any CasRx. Error bars correspond to the standard deviation for n = 4 biological replicates.

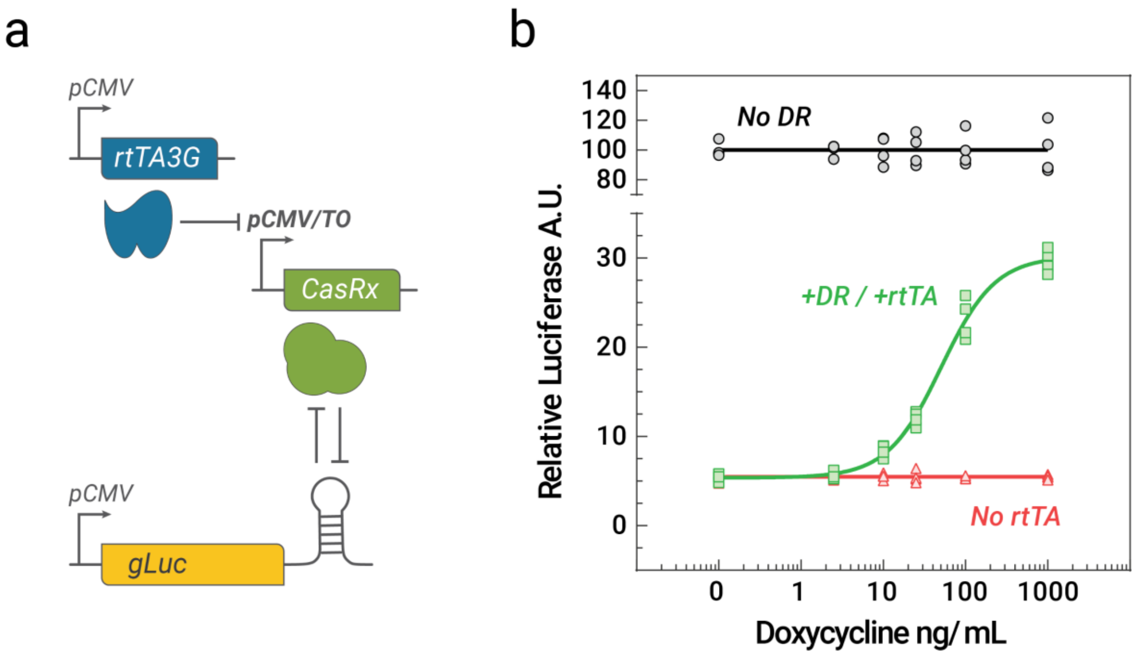


**Supplementary Figure 4. pCMV/TO-mediated repression of CasRx inhibitory function. (a)** Schematics of the experimental implementation testing pCMV/TO-mediated repression of CasRx. In the presence of doxycycline rtTA3G attaches to binding sistes downstream the pCMV/TO TATA site, causing transcriptional repression of the CasRx, that in turn should results in increasing gLuc expression **(b)** Experimental validation at the indicated concentrations of doxycycline. gLuc expression was evaluated by dual luminescence measurements resulting in the Luciferase A.U. value calculated as the ratio between gLuc and Red Firefly Luciferase luminescence for each data point. Results are plotted as Relative Luciferase A.U. representing the ratio between the Luciferase A.U. value of each data point and the mean of Luciferase A.U. values of the gLuc with DR in the absence of doxycycline. No DR (black), cells are transfected with a gLuc construct harboring no DR in the 3'UTR; +DR/+rtTA (green), cells are transfected with three plasmid encoding rtTA3G, pCMV/TO driven CasRx, and a gLuc featuring a DR in its 3'UTR; No rtTA (red), cell are not transfected with rtTA3G.

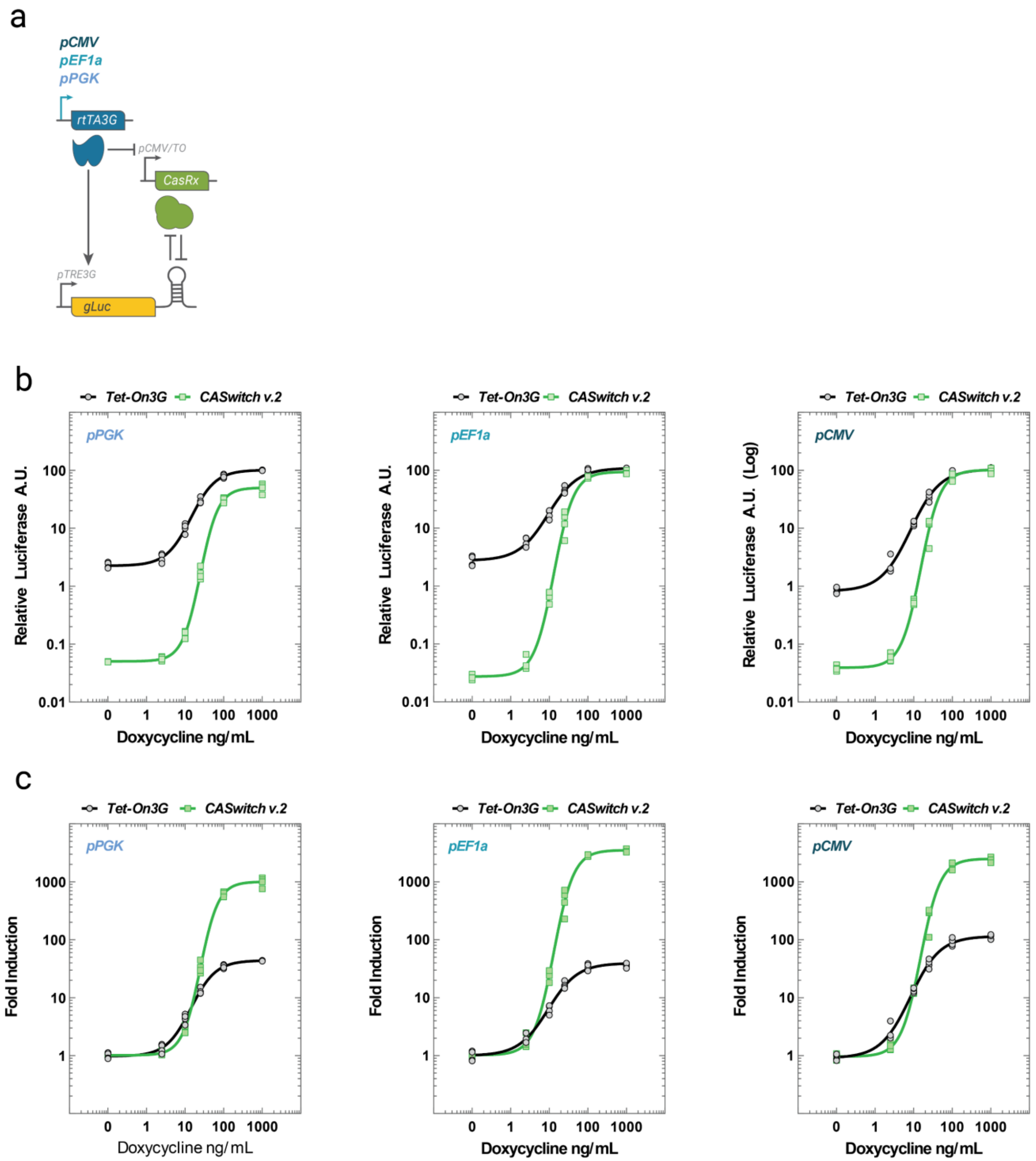


**Supplementary Figure 5. Characterization of the CASwitch v.2 system varying the expression strength of the promoter driving the rtTA3G. (a)** Schematics of the CASwitch v.2 system featuring promoters with decreasing expression strengths: pCMV, pEF1a, pPGK. **(b)** Experimental validation of CASwitch v.2 variants (green lines, promoter used indicated up left) and comparison with the state-of-the-art Tet-On3G gene expression system (black) at the indicated concentrations of doxycycline. n=4 biological replicates are shown. Relative Luciferase A.U. calculated as the percentage of the mean of Luciferase A.U. value of the Tet-On3G system at 1000 ng/mL of doxycycline and plotted in a log-scale. **(c)** Fold-induction values computed as the ratio between Relative Luciferase A.U of each data point and the mean of Relative Luciferase A.U in the absence of doxycycline.

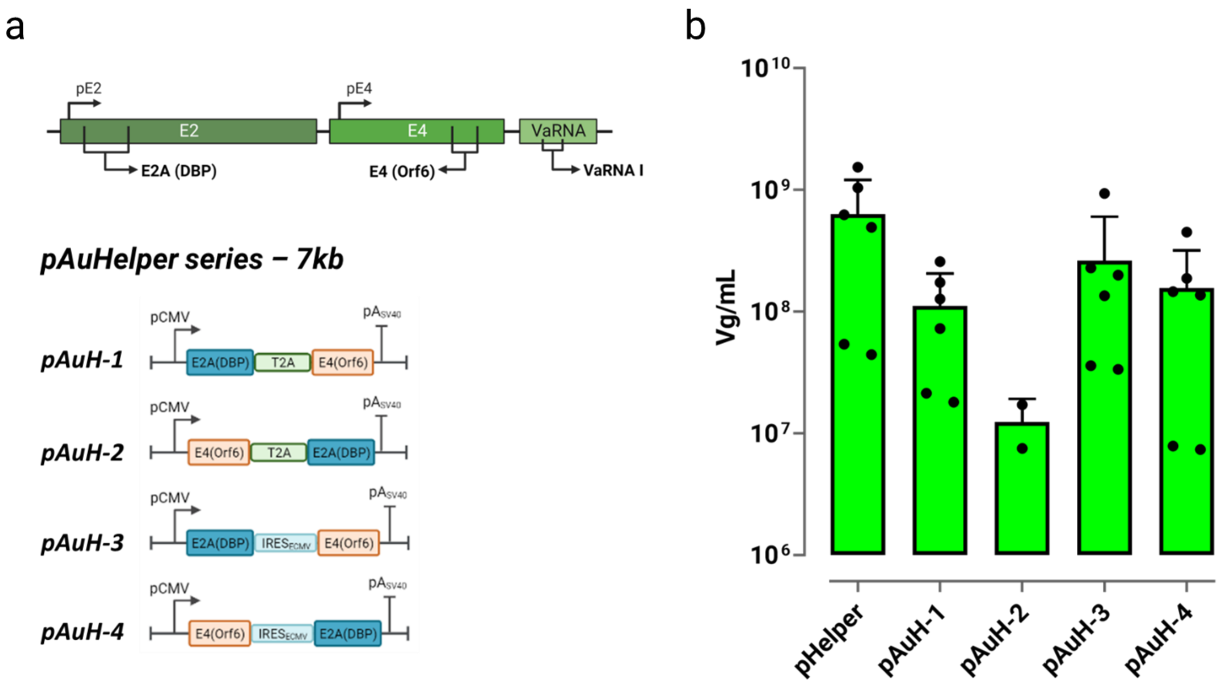


**Supplementary Figure 6. a)** Schematic of the Human Adenovirus-5 (HAdV-5) genes found in common Helper plasmids used for AAV vector manufacturing. pAµHelper plasmids schematic. E2A(DBP) and E4(Orf6) coding sequences were taken from the wt HAdV-5 genes to design a single transcriptional unit by means of the EMCV-IRES or P2A-skipping ribosome sequence. By exchanging the positions of E2A(DBP) and E4(Orf6) in the bicistronic transcriptional units, we generated four different Adenovirus Helper plasmids. **b)** AAV production yield quantification by means of qPCR. Cells were transfected with pTransgene, pPackaging, and pHelper or one of the pAµHelper plasmids. Vg/mL represents the viral genome (Vg) concentration used to quantify the production yields obtained using the indicated Helper plasmid. The error bars represent the mean and standard deviation of replicates within three independent experiments (n = 6), albeit for pAuH-2 where n = 2.

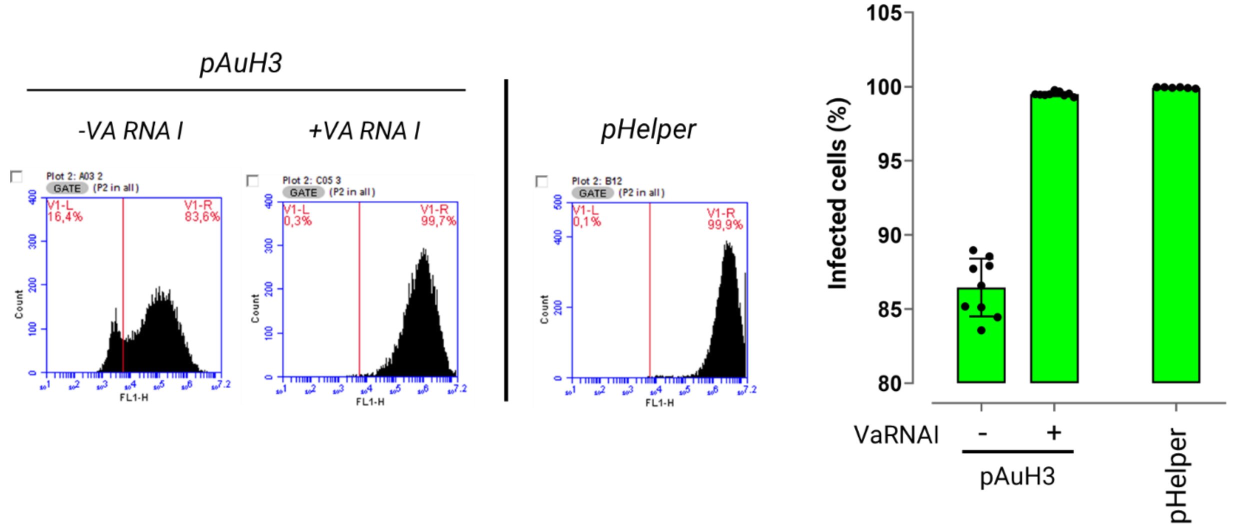


**Supplementary Figure 7. VaRNA-I fully restore production capacity for AµH-3 Helper plasmid. a)** Flow cytometry analyses of cells transduced with cell lysates containing AAV vector produced using the pAµH-3 co-transfected either with or not VaRNA-I expressing plasmid and using the standard pHelper plasmid. Representative flow cytmoterycytometry histograms (left) and their quantification (right) are also shown. The error bars represent the mean and standard deviation of n = 9 biological replicates. The percentage of transduced cells is calculated as the percentage of GFP+ cells, setting the cell autofluorescence threshold to that of non untransduced cells. At least 10,000 cells were analyzed for each point.
